## Supplementary Figures 1-10 for "Structural Basis for the Inhibition of IAPP Fibril Formation by the Hsp60 Co-Chaperonin Prefoldin"

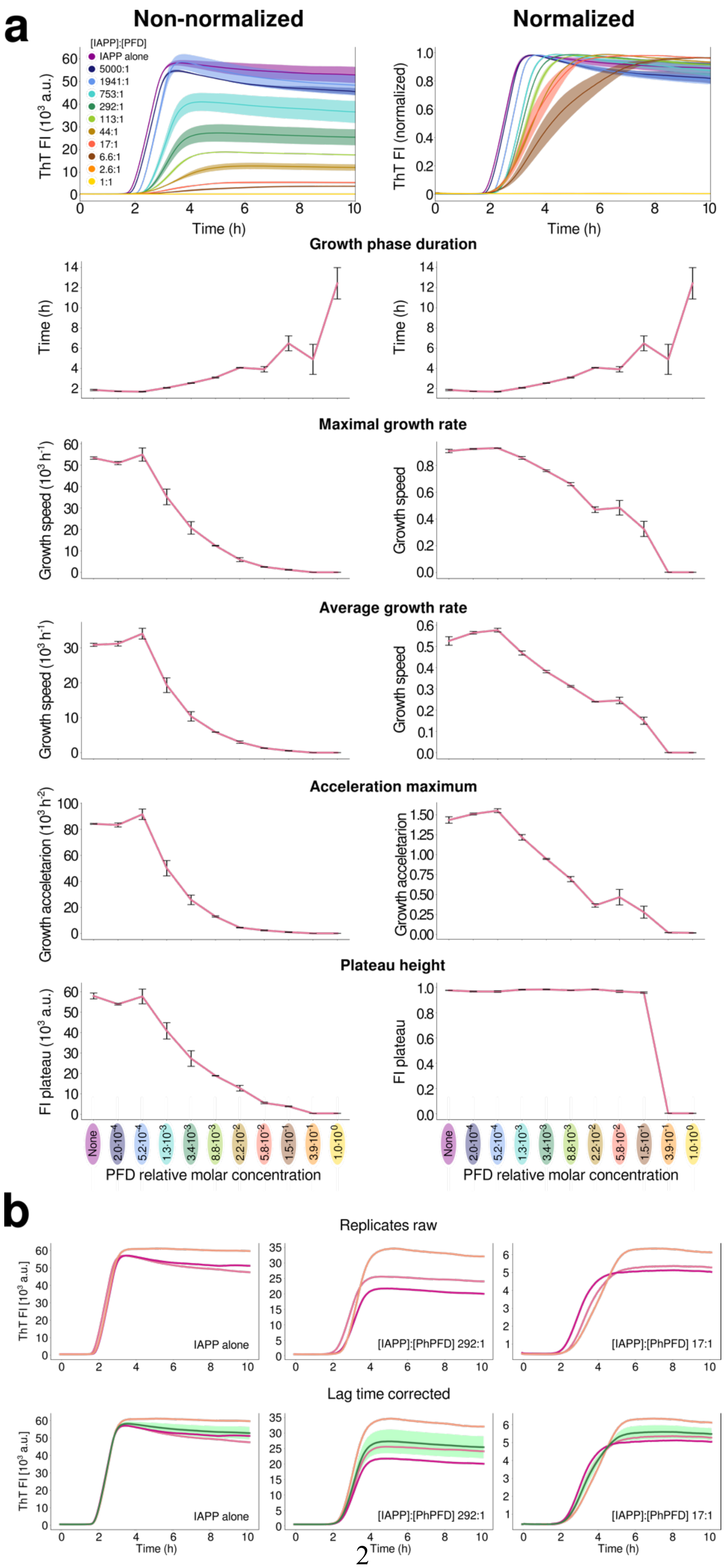
a
Non-normalized
Normalized
[IAPP]:[PFD]
IAPP alone
5000:1
1941:1
753:1
292:1
113:1
44:1
17:1
6.6:1
2.6:1
1:1
ThT FI (10³ a.u.)
ThT FI (normalized)
Time (h)
Growth phase duration
Maximal growth rate
Growth speed (10³ h⁻¹)
Growth speed
Average growth rate
Acceleration maximum
Growth acceletarion (10³ h⁻²)
Growth acceletarion
Plateau height
FI plateau (10³ a.u.)
FI plateau
None
PFD relative molar concentration
b
Replicates raw
ThT FI [10³ a.u.]
IAPP alone
[IAPP]:[PhPFD] 292:1
[IAPP]:[PhPFD] 17:1
Lag time corrected

**Supplementary Figure S1. ThT assay kinetic analysis**. **a)** Comparison of kinetic parameters obtained for non-normalized (left) and normalized (right) data. The following parameters were extracted: growth phase duration, maximal growth rate, average growth rate, acceleration maximum, and final plateau height. The analysis illustrates that the concentration dependency trends are independent of data normalization, confirming the inhibitory effect of PFD on IAPP aggregation. **b**) Exemplary demonstration of raw triplicates and their appearance after the lag time correction (averaged curves along with error shades are shown in green). The presented analysis in (a) and (b) was done on the *de novo* IAPP aggregation kinetic assay in presence of PhPFD (also shown in Fig. 1a).

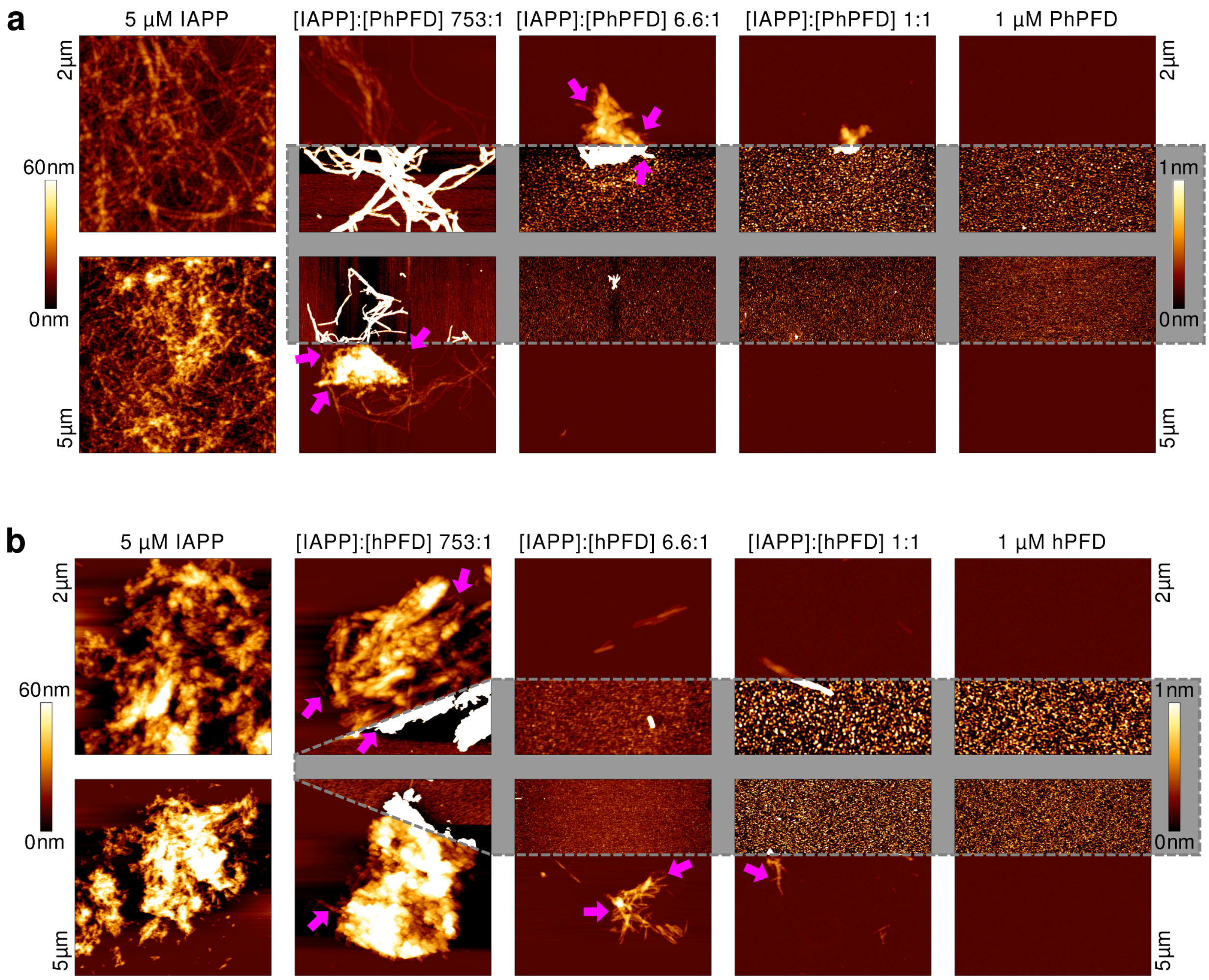


**Supplementary Figure S2. Overview of IAPP aggregate morphology in presence of PFD.** AFM images of the samples after IAPP *de novo* aggregation assays in absence and presence of different concentrations of PhPFD (a) and hPFD (b) (respective kinetics are shown in Figure 1a, top). Left to right: IAPP aggregated in absence of PFD; IAPP aggregates formed in presence of different concentrations of either PhPFD (a) or hPFD (b); prefoldin alone. The images show the close-ups of 2x2 μm$^2$ (top row) and overviews of 5x5 μm$^2$ (bottom row). For visualizing objects of diverse heights, two different colour code scaling were used: the gradient from dark brown to white represents either 0 nm to 60 nm height, or 0 nm to 1 nm (highlighted in grey). The arrows in magenta highlight individual fibrils sticking out of the bigger IAPP assemblies.

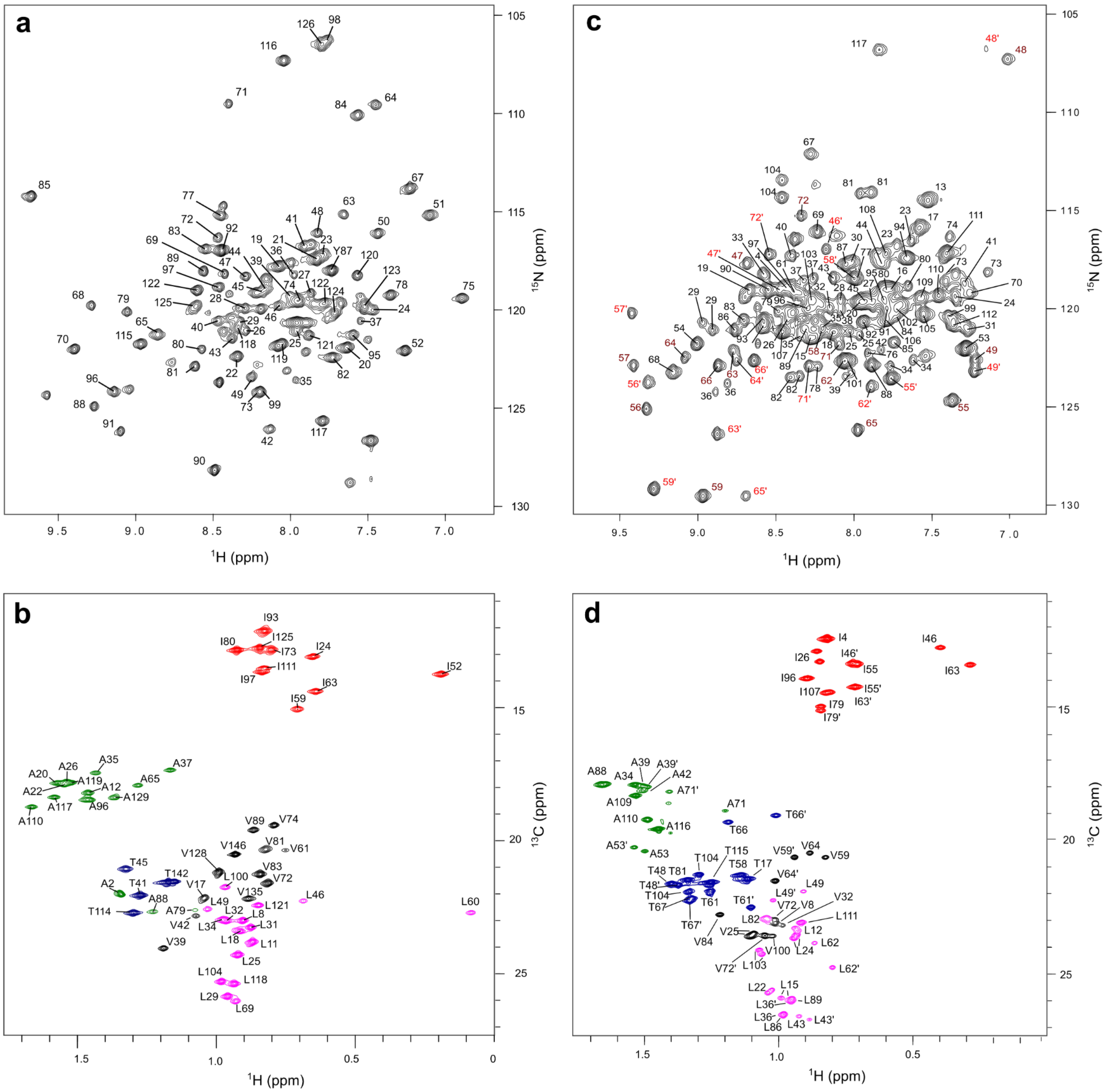


**Supplementary Figure S3. Assigned 2D $^{15}$N- and $^{13}CH_3$-TROSY spectra of PhPFD**. NMR data were recorded at 70 °C on a NMR spectrometer operating at a $^{1}$H frequency of 850 MHz. Assigned signals are annotated with corresponding residue numbers and residues which could be assigned unambiguously to β/β’ are annotated in red[38]. **a**) 2D $^{15}$N-TROSY spectrum of α−subunit. **b**) 2D $^{13}CH_3$-TROSY spectrum of α-subunit. **c**) 2D $^{15}$N-TROSY spectrum of β−subunit. **d**) 2D $^{13}CH_3$-TROSY spectrum of β-subunit.

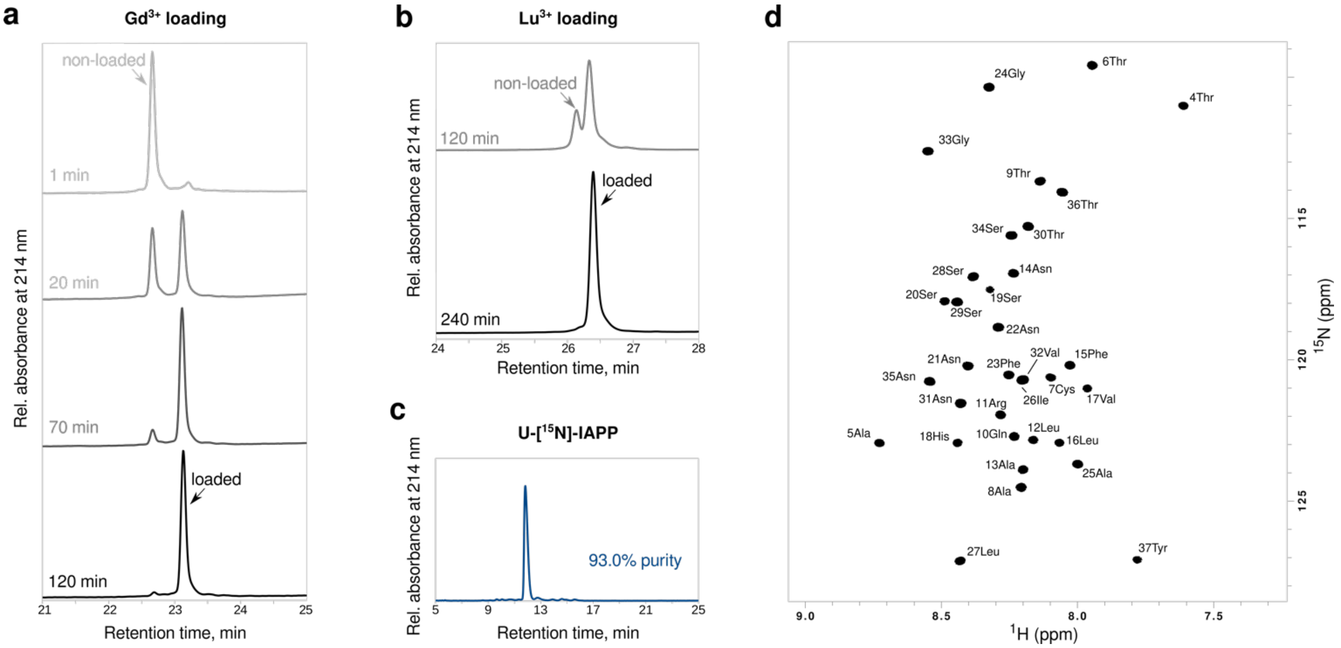


**Supplementary Figure S4. IAPP quality control.** Loading of DOTA-[β-Ala]-[β-Ala]-IAPP with $Gd^{3+}$ (**a**) or $Lu^{3+}$ (**b**) monitored by RP-HPLC at indicated incubation times. **c**) Purity of recombinant U-[$^{15}$N]-IAPP by RP-HPLC. **d**) 2D $^{15}$N-TROSY spectrum of U-[$^{15}$N]-IAPP at 10°C.

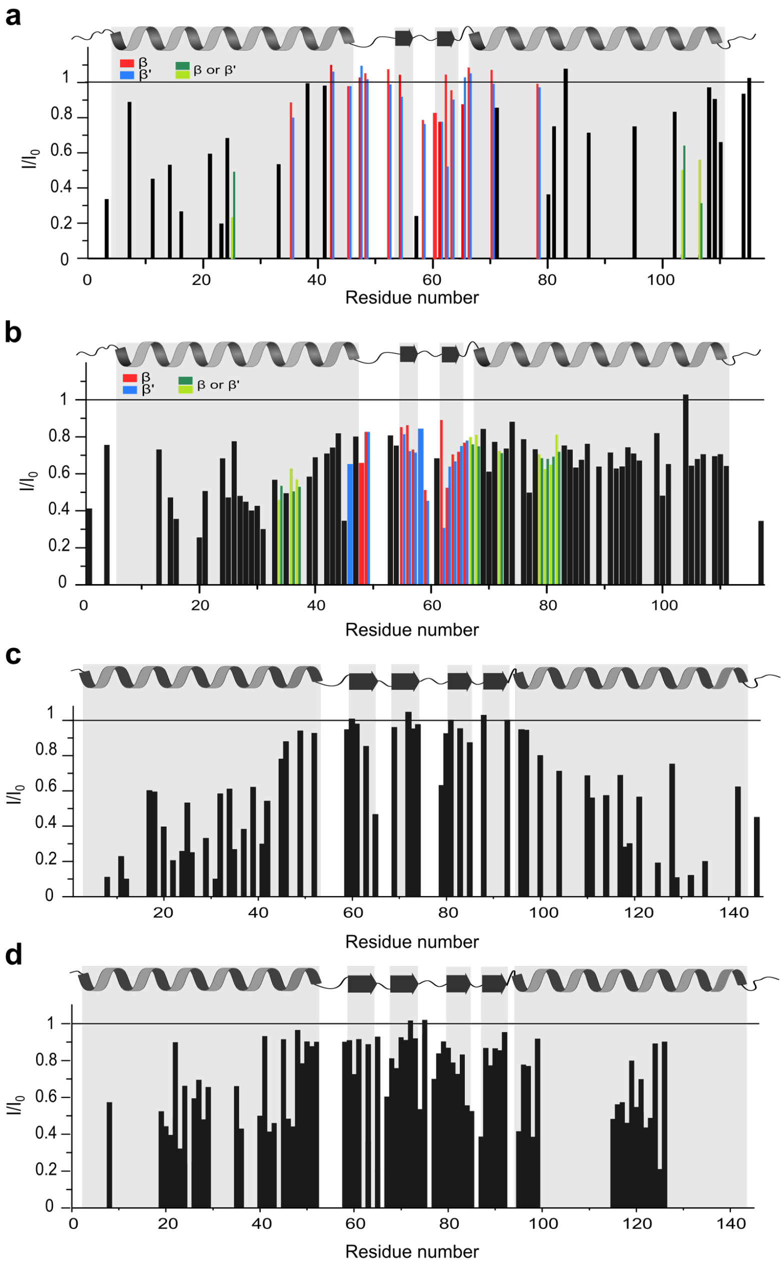


**Supplementary Figure S5. Interaction between paramagnetically labelled IAPP and PhPFD.** Histograms depicting $I_{para}/I_{dia}$ intensity ratios of PhPFD signal detected using labelled PhPFD samples in presence of Gd-IAPP or Lu-IAPP, respectively. Reconstituted PhPFD with β subunits (a,b) and α subunits (c,d) labelled on $^{13}CH_3$ groups (a,c) or $^{15}$NH-backbone groups (b,d) were mixed with Lu/Gd-loaded IAPP in a ratio of 1:2 ($^{13}CH_3$-labeling) or 1:1 ($^{15}$N-labeling). No values indicate missing assignment[38] or absence of methyl-groups.

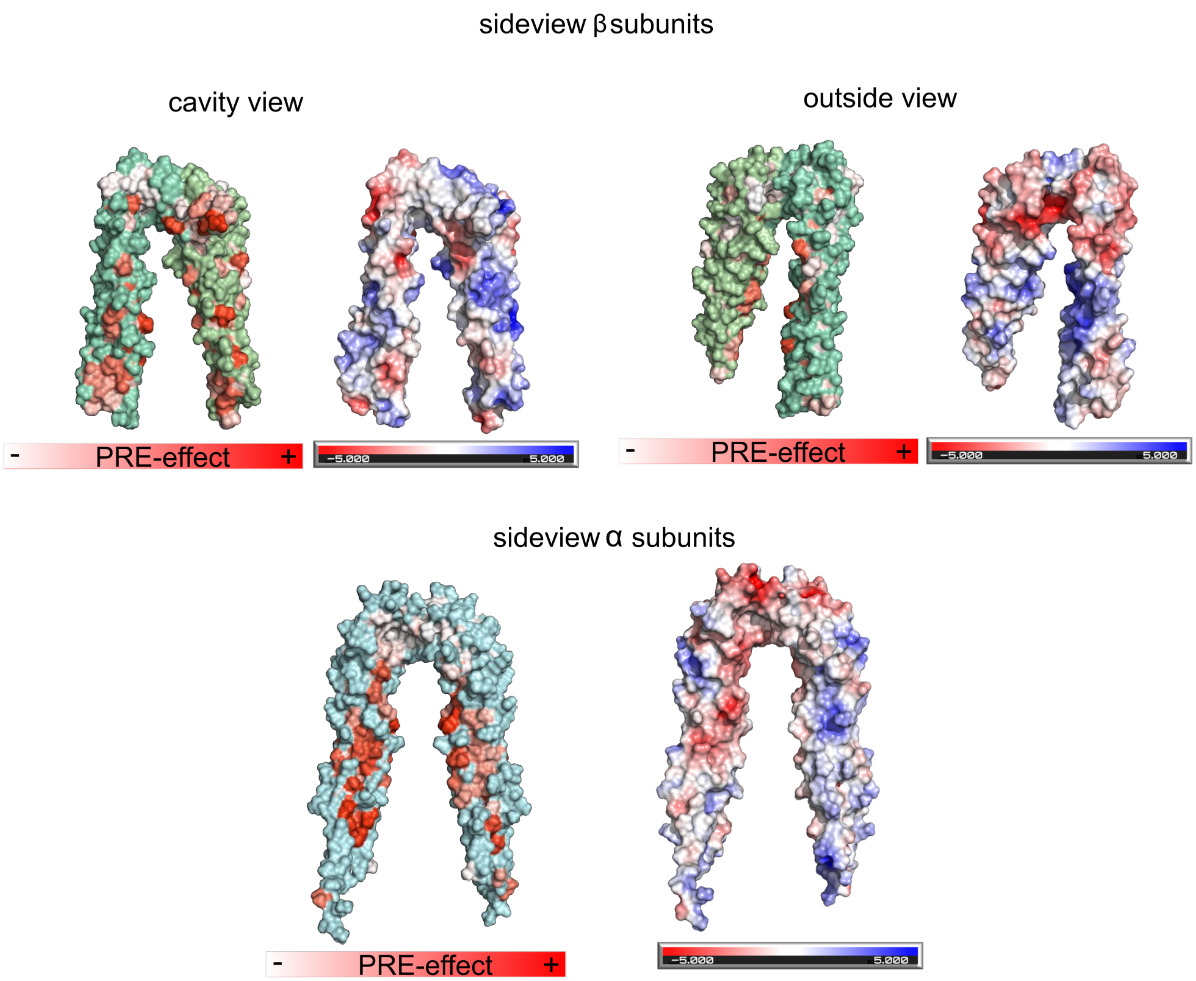


**Supplementary Figure S6. Comparison of PRE-mapping and electrostatic surface on PhPFD.** The interaction surface mapped on PhPFD by paramagnetic relaxation enhancement juxtaposed with the theoretical electrostatic surface computed by APBS electrostatics implemented in pymol[73].

**a**

| | Vector (antibiotic resistance) | Sequence |
|---|---|---|
| **PFD1 (beta)**<br>**14.2 kDa** | **pET-21a**<br>**(Ampicillin)** | MAAPVDLELK KAFTELQAKV IDTQQKVKLA DIQIEQLNRT KKHAHLTDTE IMTLVDETNM YEGVGRMFIL QSKEAIHSQL LEKQKIAEEK IKELEQKKSY LERSVKEAED NIREMLMARR AQ |
| **PFD2 (beta)**<br>**16.9 kDa**<br>**with tag 18.8 kDa** | **pET-41**<br>**(Kanamycin)** | MGSSHHHHHH SSGLVPRGSH MAENSGRAGK SSGSGAGKGA VSAEQVIAGF NRLRQEQRGL ASKAAELEME LNEHSLVIDT LKEVDETRKC YRMVGGVLVE RTVKEVLPAL ENNKEQIQKI IETLTQQLQA KGKELNEFRE KHNIRLMGED EKPAAKENSE GAGAKASSAG VLVS |
| **PFD3 (alpha)**<br>**18.7 kDa** | **pET-21a**<br>**(Ampicilin)** | MKQPGNETAD TVLKKLDEQY QKYKFMELNL AQKKRRLKGQ IPEIKQTLEI LKYMQKKKES TNSMETRFLL ADNLYCKASV PPTDKVCLWL GANVMLEYDI DEAQALLEKN LSTATKNLDS LEEDLDFLRD QFTTTEVNMA RVYNWDVKRR NKDDSTKNKA |
| **PFD4 (beta)**<br>**15.2 kDa**<br>**with tag 17.1 kDa** | **PET-41**<br>**(Kanamycin)** | MGSSHHHHHH SSGLVPRGSH MKKAAAEDVN VTFEDQQKIN KFARNTSRIT ELKEEIEVKK KQLQNLEDAC DDIMLADDDC LMIPYQIGDV FISHSQEETQ EMLEEAKKNL QEEIDALESR VESIQRVLAD LKVQLYAKFG SNINLEADES |
| **PFD5 (alpha)**<br>**17.3 kDa** | **pET-21a**<br>**(Ampicillin)** | MAQSINITEL NLPQLEMLKN QLDQEVEFLS TSIAQLKVVQ TKYVEAKDCL NVLNKSNEGK ELLVPLTSSM YVPGKLHDVE HVLIDVGTGY YVEKTAEDAK DFFKRKIDFL TKQMEKIQPA LQEKHAMKQA VMEMMSQKIQ QLTALGAAQA TAKA |
| **PFD6 (beta)**<br>**14.6 kDa** | **pET-21a**<br>**(Ampicillin)** | MAELIQKKLQ GEVEKYQQLQ KDLSKSMSGR QKLEAQLTEN NIVKEELALL DGSNVVFKLL GPVLVKQELG EARATVGKRL DYITAEIKRY ESQLRDLERQ SEQQRETLAQ LQQEFQRAQA AKAGAPGKA |

**b**

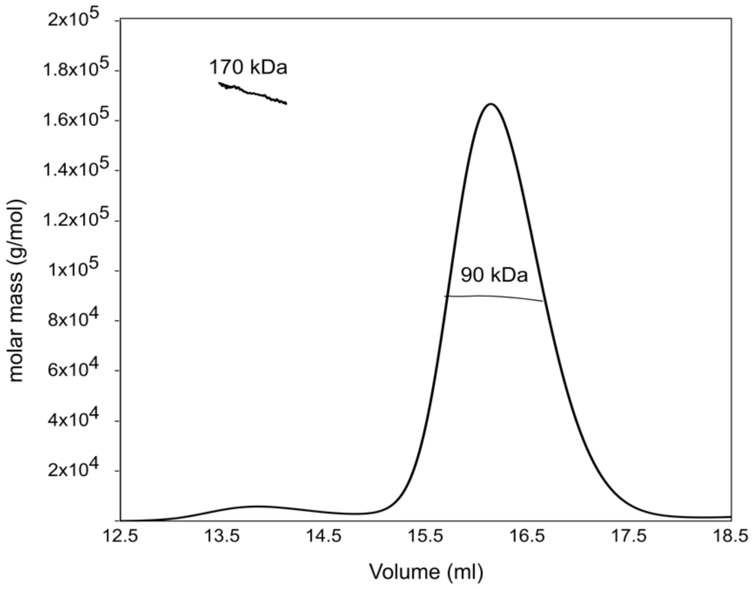


**c**

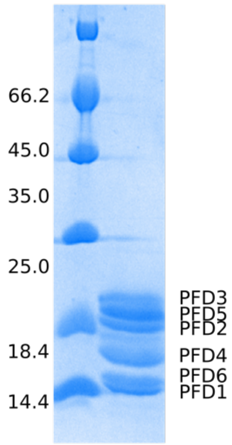


**Supplementary Figure S7. Production of hPFD samples and quality control**. **a**) Summary of information on hPFD subunit sequences (PFD2 and PFD4 include polyhistidine tag sequences) and plasmids used in this study. **b**) SEC-MALS analysis of heterohexameric hPFD sample. SEC-MALS analysis showed two species, a minor species at about 170 kDa (~5%) and a major one at about 90 kDa (~95%). The formation of two species, the hexameric complex and a complex of double size was already observed by Aikawa et al.[54] **c**) SDS-PAGE of sample after purification. On the SDS-PAGE six bands appear, confirming the formation of a pure hPFD complex.

**a**

| | Vector (antibiotic resistance) | Sequence |
|---|---|---|
| **β-subunit 13.3 kDA** | **pET23c (Ampicillin)** | MQNIPPQVQA MLGQLDTYQQ QLQLVIQQKQ KVQADLNEAK KALEEIETLP DDAQIYKTVG TLIVKTTKEK AVQELKEKI ETLEVRLNAL NRQEQKINEK VKELTQKIQA ALRPPTAG |
| **α-subunit 16.6 kDa** | **pET23c (Ampicillin)** | MAQNNKELEK LAYEYQVLQA QAQILAQNLE LLNLAKAEVQ TVRETLENLK KIEEEKPEIL VPIGAGSFLK GVIVDKNNAI VSVGSGYAVE RSIDEAIGFL EKRLKEYDEA IKKTQGALAE LEKRIGEVAR KAQEVQQKQS MTSFKVKK |

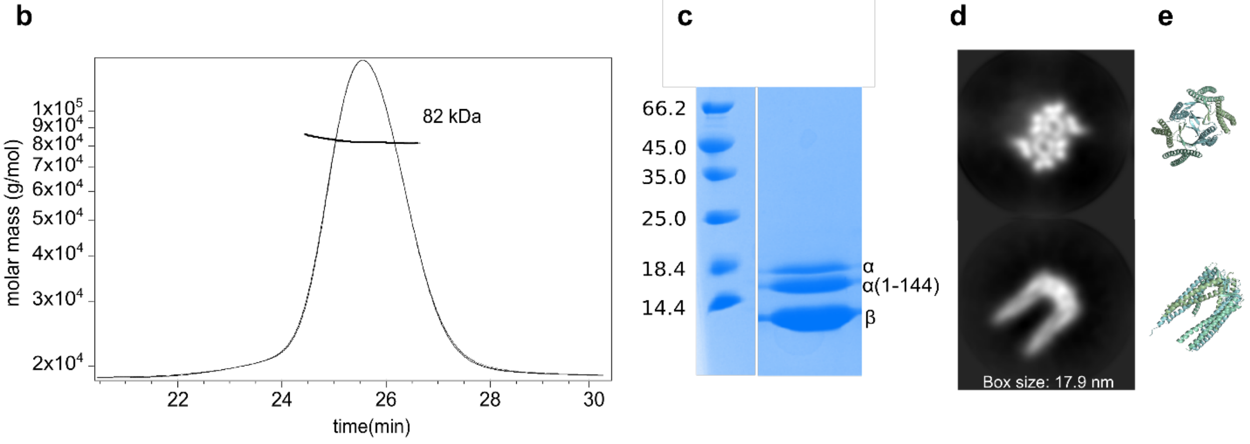


**Supplementary Figure S8. PhPFD production and quality control. a**) Summary of information on PhPFD α and β subunits sequences and plasmids used in this study. **b**) SEC-MALS analysis of $\alpha_2\beta_4$–PhPFD sample. The mass of PhPFD was estimated to be 82 kDa. **c**) SDS-PAGE analysis and subsequent mass spectrometry analysis (not shown) indicated the presence of a partial cleavage on the C-terminal residues 145-148 of the α-subunit. This cleavage located in the unstructured C-terminal sequence does not hinder the formation of the complex, as can be observed from (**d**) the 2D classes obtained from cryo-EM analysis. Formation of the β-barrel structures with the six pairs of protruding helices can be clearly seen from the top view. Formation of the β-coiled coil α-helices is also observed in the side view. The other side view is missing due to preferential orientation hampering determination of 3D structures from cryo-EM images. In (**e**) the according views of the molecular model are shown.

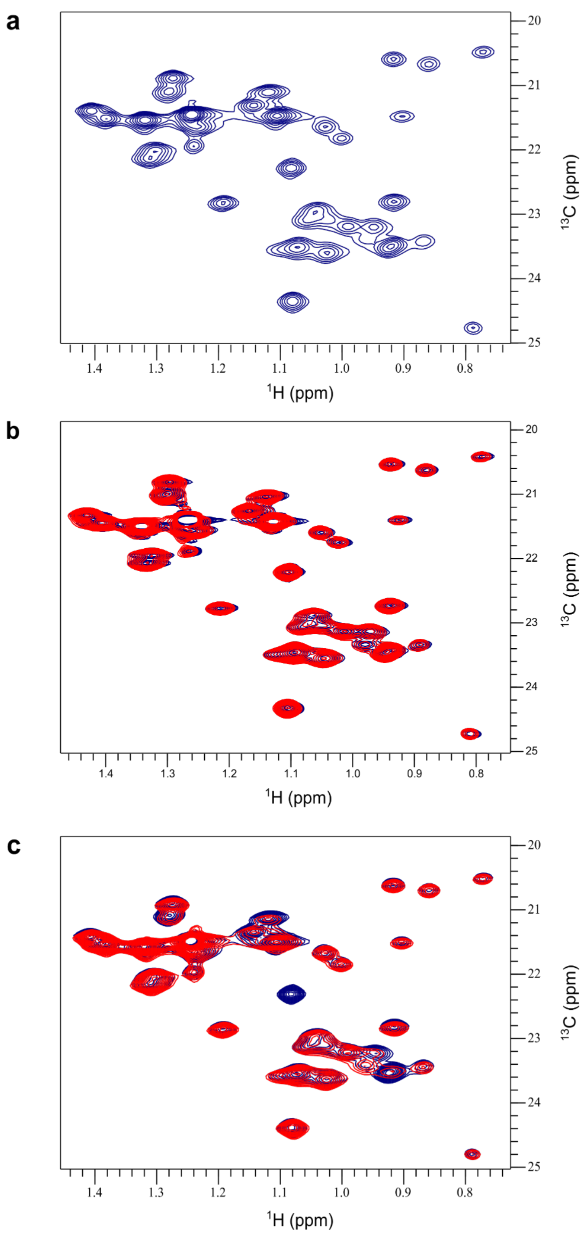


**Supplementary Figure S9. Interaction between paramagnetically labelled IAPP and PhPFD**. **a**) Reference $^{13}CH_3$-TROSY spectrum of PhPFD labelled on methyl residues on the β subunits at 30 °C in 25mM MES/NaOH (pH 6.5), 25mM $MgCl_2$. **b**) PhPFD control spectra in presence of 1:1 of $Lu^{3+}$-loaded DOTA (blue) or $Gd^{3+}$-loaded DOTA (red). **c**) PRE effect detected on $^{13}CH_3$-labelled PhPFD in presence of diamagnetic Lu-IAPP (blue) or paramagnetic Gd-IAPP (red).

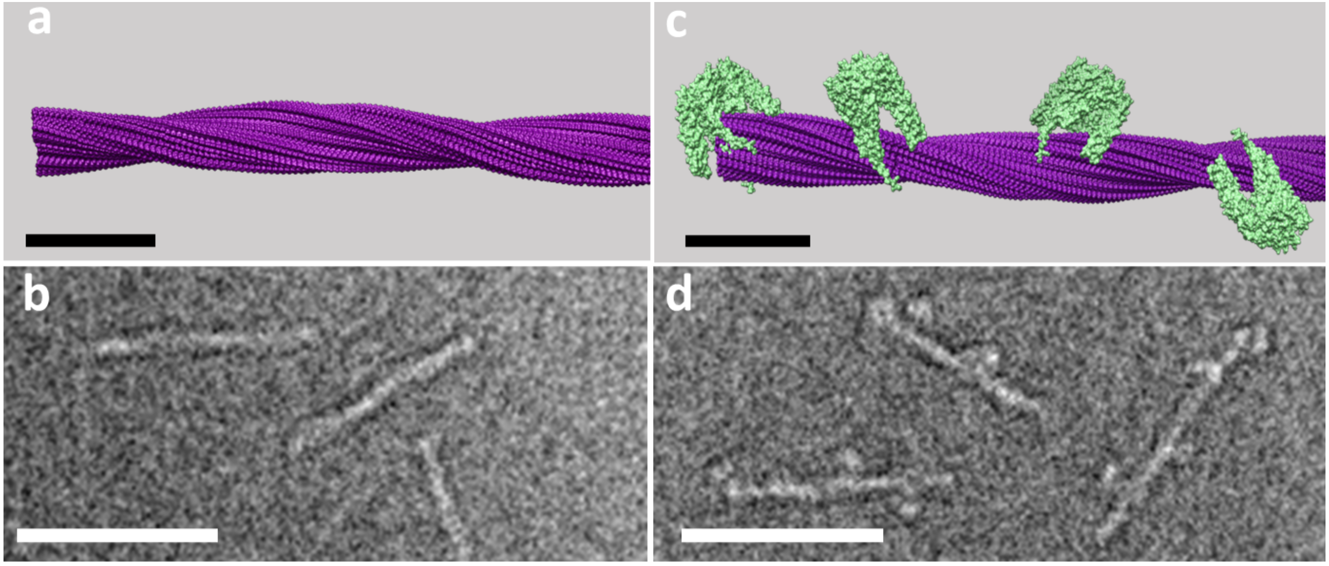


**Supplementary Figure S10. Simulated EM images of PhPFD bound to IAPP fibrils.** Transmission electron microscopy images were simulated with TEM-simulator[74] (http://tem-simulator.sourceforge.net/), using the model shown in Fig. 4i. **a**) Structure of the ordered part (residues 13-37) of polymorph 1 IAPP fibril. **b**) Simulated TEM image of IAPP fibrils according to the structure depicted in (a). **c**) Model of interaction between IAPP fibril and PhPFD. **d**) Simulated TEM image of IAPP fibril decorated with PhPFD. Horizontal black and white bars correspond to a length of 10 nm and 50 nm, respectively.
